# Adult Zebrafish Engage in Path Integration-like Behavior When Exploring a Novel Environment

**DOI:** 10.64898/2025.12.31.697215

**Authors:** Bhavjeet Sanghera, Philip Shraybman, Jagmeet S. Kanwal

## Abstract

Spatial navigation relies on the integration of self-motion (idiothetic) and external (allothetic) cues to construct an internal representation of space. While path integration has been extensively studied in insects and mammals, its presence and mechanistic basis in non-mammalian vertebrates remain poorly understood. Here, we show that adult zebrafish (*Danio rerio*) placed in a novel, visually impoverished environment exhibit systematic reorganization of exploratory behavior consistent with path integration-like navigation. Using markerless tracking, we identified a sequence of stereotyped bouts initiated by wall contact that consists of subsequent reorientations followed by a corrective sharp turn, returning the animal toward its initial contact location. Markov chain analyses reveal structured transitions linking wall-touch events, reorientation states, and high-curvature turns early in exploration, followed by a progressive decoupling of turning from boundary contact over time. Angular error-correction analyses demonstrate that cumulative reorientation predicts the magnitude and direction of subsequent sharp turns, consistent with vector-based error compensation. Multivariate analyses further show a graded shift in behavioral state space from early to late exploration, rather than an abrupt strategy switch. Together, these results provide evidence that zebrafish employ a go-and-touch strategy that integrates tactile boundary information with self-motion cues to estimate spatial layout. Our findings establish zebrafish as a vertebrate model for studying the computational and neurogenetic basis of path integration that develops by the fifth week post-fertilization.

## 1. Introduction

Having knowledge of one’s immediate environment is crucial for survival of any species. The process of gaining this knowledge is initiated as soon as an organism can locomote in its environment. For organisms to successfully navigate within their environment, they need to either construct a mental representation of their home space using idiothetic information or possess a mechanism of using allothetic cues as stable reference or waypoints (Vickerstaff and Di Paolo, 2005; Whishaw and Brooks, 1999). These cues can include polarized light, magnetic fields, celestial objects or landmarks in the terrestrial environment (Fleischmann et al., 2018; Freas et al., 2024; Grob et al., 2024; Lebhardt and Ronacher, 2014; Legge et al., 2014). A reliable and systematic representation of the home space allows the organism to find food, mates for reproduction, and rest or shelter demanded by adverse weather conditions. During exploratory excursions out of their home range, animals also need to escape unknown territory when threatened by a predator or a conspecific and quickly return to their shelter (Ellison et al., 2020; Giuggioli and Kenkre, 2014). Navigational brain maps can save energy by providing a knowledge base to compute a direct path back to the home coordinates (de Cothi et al., 2022; Kanwal, 2024; Teng et al., 2015; Tsoar et al., 2011). The phenomenon of computing a short path back has been termed as "path integration" (Conklin and Eliasmith, 2005; Wehner and Srinivasan, 1981). Path integration (PI) may be considered as a fundamental navigation process by which an organism calculates its position relative to a starting point by continuously updating its position and orientation based on self-movement and environmental cues (Etienne et al., 2004). The direction and distance traveled are key idiothetic cues, although speed, and time taken to travel between waypoints can also play an important role in computing the shortest return path (Knierim et al., 1998; Lee et al., 2013; Whishaw and Brooks, 1999). PI can be a mechanism to save time and energy for foraging as well as a mechanism for confirmation of a cognitive map and reinforcement of knowledge of one’s environment (Kanwal, 2024).

Taking advantage of sensory cues and motor abilities, a newborn of any species can begin to construct a spatial representation of its environmental space within its neural space (Langston et al., 2010). How this happens has been a focus of intense debate over the last decade and has led to significant scientific breakthroughs in our understanding of brain-based navigational systems (Donoghue et al., 2023; Kaske et al., 2006; Moser et al., 2008; Rolls, 2021; Vinepinsky et al., 2020). Much of the new understanding has come from studies in rats, but recent studies in other species, such as bats and primates (including humans) are helping to advance our understanding of interactions between key brain regions and the spatial receptive field properties of specialized cells, such as head-direction, place- grid- and border cells that play key roles in enabling navigation (Danilovich and Yovel, 2019; Iggena et al., 2023; Knierim et al., 1998; Menti et al., 2022; Moser et al., 2008). Some of these cell types were recently demonstrated to be present within the telencephalon of goldfish (Saito and Watanabe, 2006; Vinepinsky et al., 2020). In fish, these cells are present in an area homologous to the hippocampus of mammals. Following our initial observation of PI-like behavior in adult zebrafish, place cells were also reported in the telencephalon of larval zebrafish (Yang et al., 2024). Taken together, these findings support the possibility that, similar to mammals, fish may use cognitive maps for spatial navigation without well organized hippocampal and entorhinal-like well organized brain structures.

Zebrafish are highly visual and typically use visual cues for navigating in their habitat, though they can also thrive in turbid waters (Sundin et al., 2019). We assumed that zebrafish use allothetic cues, such as stable objects and landmarks in their aquatic environment to map their swim-trajectories within their home space, which includes slow-running streams and shallow ponds exposed to bright light (Sundin et al., 2019). Changing water levels during the monsoon rains, however, can rapidly alter the underwater scenery. Fish need to continue to navigate their surroundings for foraging, predator escape and homing during these changing conditions as well as when the waters become murky. This is possible only if they can rapidly re-map their environment by attending to and integrating new sensory cues with idiothetic information.

Considering the rapid changes that can occur within their natural habitat, here we test the hypothesis that when adult zebrafish are placed within a novel environment, they indulge in a systematic exploration of their new environment. Depriving the environment of clear under- and above-water visual landmarks may unmask systematic patterns of exploration behavior used in natural environments. At a minimum, their free-swimming behavior may undergo subtle yet definitive changes over time as they begin to remap their environment. Alternately, zebrafish may simply exhibit small random deviations in stereotypic swim patterns determined largely by established self-motion bodily (idiothetic) cues. As a test, we define and examine changes in postural and locomotory patterns of free-swimming behavior. If swim patterns shift over time, we were curious if zebrafish adop a particular strategy to explore and cognitively re-map their new environment.

How animals navigate novel environments without reliable visual landmarks remains a central question in neuroscience. We show that adult zebrafish reorganize exploratory behavior over time, using boundary contact to scaffold internal representations of space and guide navigation through a path integration–like strategy. By revealing structured behavioral algorithms and statistically robust state-transition dynamics in a genetically tractable vertebrate, this work establishes zebrafish as a powerful model for uncovering the neural and evolutionary foundations of spatial navigation. We discuss our findings together with what is known about neurobehavioral mechanisms for spatial navigation and map-making strategies reported in other species.

## 2. Methods

### 2.1. Animal acquisition and maintenanced

Adult (6-12 months old) wild-type zebrafish (*Danio rerio*) were used in this study. Fish were housed in groups of 10 fish per 2.5-liter tank which were filled with continuously aerated and filtered habitat water and kept at 26°C in a 14:10 light:dark cycle in the zebrafish facility at Georgetown University Medical Center, Washington, DC. They were fed daily with specially designed dry flake food. A total of nine adult zebrafish and nine larvae (7 male; 2 female) were used for recordings. Prior to experiments, fish were transferred from their home tank in the zebrafish facility to a separate 2.5-liter holding tank containing habitat water and transported to the laboratory. To minimize stress from out-of-water handling, fish were transferred between tanks using a small plastic scoop so that fish were always contained within a small volume of habitat water. All procedures were approved by the Institutional Animal Care and Use Committee.

### 2.2. Recording apparatus

Fish transferred to the holding tank were allowed to acclimate to the environmental conditions of the test environment. Fish were scooped out once again from the holding tank and introduced to the center of the small, oval test tank (11 cm by 17 cm) through an opaque plastic funnel. The test-tank had a smooth, curved and opaque, milky-white ceramic surface. It was illuminated with diffuse light that produced uniform illumination over the tank and water surface (Fig. 1A). It is important to note that the gentle curve and sloping wall of the tank wall did not introduce any corners, and the tank bottom was visually continuous with the sides, eliminating discernable tank boundaries. A document camera (Okiocam, Okiolabs, Inc.) mounted 35 cm above the water surface was used to continuously record free-swimming behavior of the zebrafish at a resolution of 1024 by 768 p. The 3 cm by 4 cm camera face was the only well-defined overhead object visible to the fish from below. Water level was maintained at ∼ 2.0 cm to keep fish in the plane of focus for two-dimensional tracking and position analysis. Video recordings were obtained for 10 minutes, at 30 fps (frames/s) using video capture software (Okiocam recorder, vers. 1.03; Okiolabs, Inc.), immediately after the fish were introduced into the tank. All data were saved on the hard drive of an Apple MacBook Pro computer and transferred to Dropbox for backup storage and data access and analysis using a Microsoft Windows computer equipped with a GeForce NVIDIA Graphics Processor Unit (GPU) card for further analysis.

**Figure 1.**
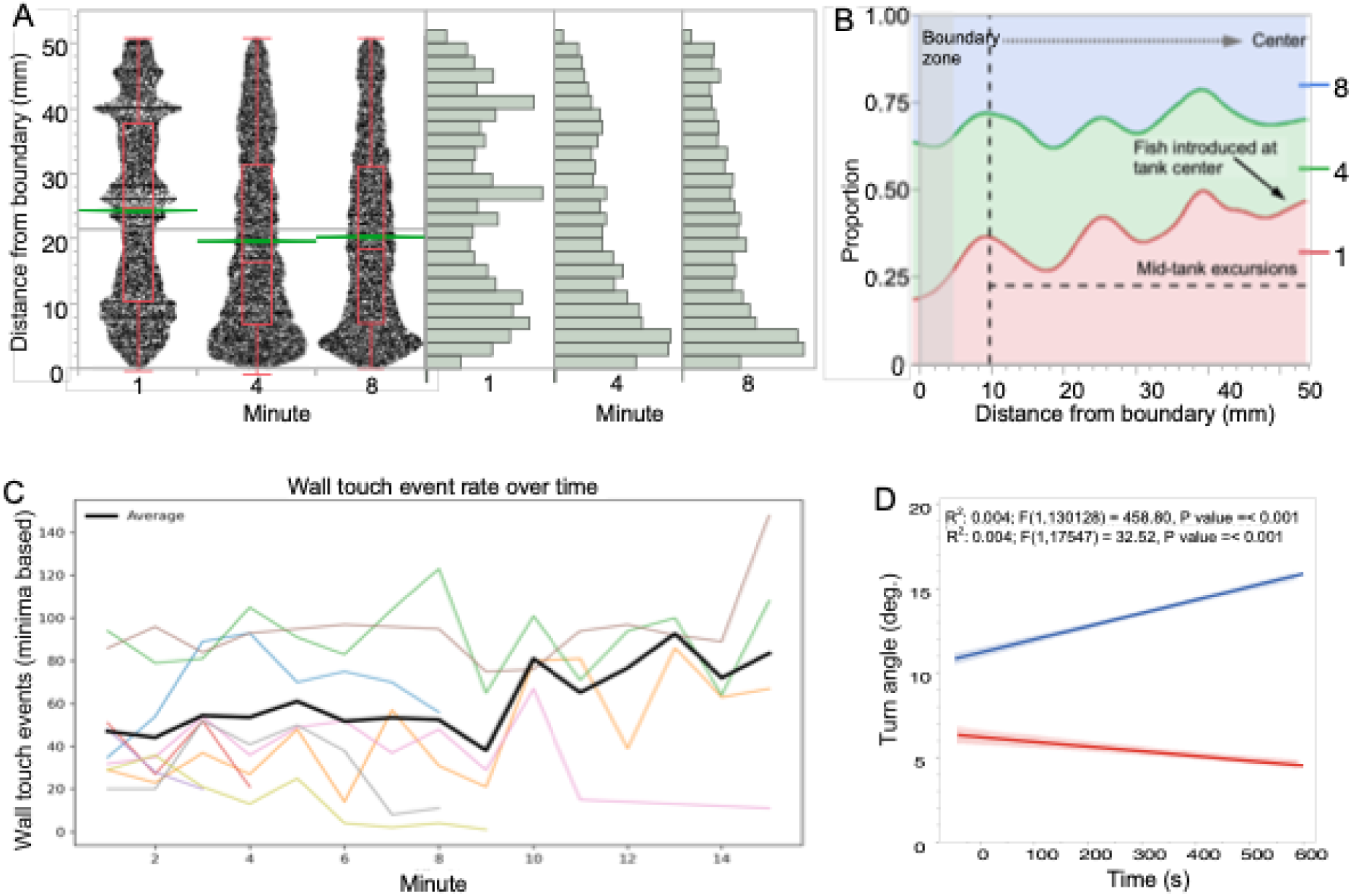
A. Box plots superimposed on scatterplots of distance-to-boundary of tank in first and last minute of recording averaged across all 9 animals. Histograms on the right show the distribution of distances in both instances. Distances were measured between the ocular center and the tank boundary. The smaller median distance in the tenth minute indicate that fish spent the bulk of their time along the sides of the tank. B. Density plots of fish distance from the center for the first (red), fifth (green) and the tenth (blue) minute of recording. The data show parallel waves in space, indicating regions during each minute where fish spent proportionately more time (long distances of OC from the boundary) and a plateau for locations in the mid-region. Region identified as wall touch is indicated by gray zone and color coding of minutes is indicated on the right. C. Line plots showing the progression of wall touch events over time. D. Linear fit lines for body angles computed from head transitions over time for locations near tank boundary (red) and within mid-regions of the tank (blue). Shaded zones show a 90% confidence interval.

### 2.3. Animal tracking and data analysis pipeline

We used the DeepLabCut (DLC) software to train a markerless pose-estimation model to extract x-y coordinates from up to eight body parts along the longitudinal axis of the fish (Figure 1B). The x-y coordinates of the midpoint of the head between the two eyes labeled as the “ocular-center” (OC), “mid-flank (MF)”, “tailfin-base”, and “tailfin-midpoint” were used as markers to extract and identify body configurations. Frame-by-frame coordinates for each marker in the x-y plane were saved in a spreadsheet. From these coordinates, the body displacement, body posture (angular flexion of body at midflank), heading (orientation of straight line between OC and MF) and distance from tank boundary were calculated. We leveraged B-SoiD (Behavioral Segmentation of Open-field in DeepLabCut) and SimBA (Simple Behavioral Analysis), software packages implementing a machine learning algorithm for pose estimation, to visualize possible body postures and action patterns that adult zebrafish engage in (Hsu and Yttri, 2021; Mathis et al., 2018; Nilsson et al., 2020). B-SOiD calculates features like distance between markers, angles, and speed before performing dimension reduction for cluster analysis as depicted in (Fig. S1). We previously developed a classification scheme from these data for categorizing, identifying and naming zebrafish postures and movement patterns in free-swimming adult zebrafish ((Kanwal et al., 2025); Sanghera and Kanwal, in preparation). Translational movements were tracked using head (OC) coordinates. Per our analysis pipeline, we extracted swim and turn parameters that included forward-swim, stationarity, reorientation, and sharp-turn (Fig. S2). We computed the probability of state (pose and action) transitions to determine the behavior patterns exhibited by each individual during exploration of the novel tank environment.

### 2.4. Statistical considerations and justification

#### 2.4.1. Geometric analysis and behavioral metrics

We used JMPpro (vers. 8), the statistical package R as well as custom-generated Python scripts for data analysis. Because our primary goal was to characterize structure and trajectories of boundary interaction strategies rather than to test a single population-level hypothesis, we adopted an event-centric and within-animal analytical framework. Wall-contact (wall touch or WT) behavior was quantified using minima-based events, avoiding inflation due to prolonged contact and ensuring that values used to generate the probabilistic network corresponded to discrete behavioral bouts.

Spatial positions of events were projected onto a one-dimensional boundary coordinate (*s*) representing cumulative arc length along the tank perimeter, estimated using a capsule approximation of tank geometry. For each fish, *s* was phase-aligned such that the first wall-contact event in the session occurred at *s* = 0, enabling comparison of revisitation structure across animals. Circular statistics were computed on aligned event positions, including normalized circular entropy (quantifying spatial dispersion), median absolute deviation (MAD) of |*s*| (quantifying alignment stability), and return probability (fraction of events within |*s*| < 20 mm of the aligned origin). These analyses are specifically designed to reveal within-subject organization and stability that may be obscured by group-mean statistics, particularly in small cohorts. Comparisons between early (Minutes 1–2) and late (Minutes 7–8) periods were performed using paired, non-parametric Wilcoxon signed-rank tests, appropriate for small sample sizes and non-Gaussian distributions (see Table S1).

#### 2.4.2. Markov transition analysis

For each fish and minute, a first-order Markov transition matrix was constructed from consecutive frame-wise state labels. Transition counts were normalized row-wise to yield conditional probabilities. To prevent dominance by highly active individuals, group-level matrices were computed as the mean of per-fish transition matrices, ensuring equal weighting across animals. Markov networks (network plots based on transition matrices that compute the frequency of occurrence of postural configurations and their transitions) were visualized with node size proportional to state occupancy (fraction of frames). Edges or rays between nodes were used to depict a probability distribution, p(x), with line/arrow thickness proportional to transition probability. Node placement was determined by spring layouts weighted by state co-occurrence frequency and arrowheads specified directionality of reciprocal transitions.

To quantify experience-dependent changes, we computed difference networks (ΔP) as:

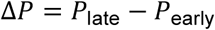

where early and late correspond to Minute 1 vs Minute 8. Edges in difference networks were color-coded (red = increased, blue = decreased) and annotated with signed Δ values.

#### 2.4.3. Quantification of path integration–like navigation from WT-anchored bouts

To test whether zebrafish exhibit path integration–like navigation, we analyzed WT-anchored navigation bouts and quantified return specificity and vector-based home alignment using self-motion cues. WT events were identified using a conservative geometric criterion (distance to tank wall < 4mm). For each fish, WT onsets separated by at least *N* frames (to ensure independence) were treated as anchor events, defining an anchor position 𝐴 = (𝑥_0_, 𝑦_0_) and an anchor boundary coordinate 𝑠_0_(arc-length parameterization along the rounded tank boundary).

For each anchor event, we defined a navigation bout beginning at the anchor frame and extending until the next boundary contact or the end of the minute. The “outbound phase” was defined as frames following the anchor during which the fish moved away from the boundary (distance to boundary exceeding a threshold for ≥M consecutive frames). The “return phase” was defined as frames during which the fish either decreased its distance to the anchor or re-contacted the boundary. Bouts lacking a clear outbound and return phase were excluded.

#### 2.4.4. Vector-based home alignment and experience-dependent changes

To test whether fish aligned their heading toward a predicted home direction, we computed the integrated displacement vector from the anchor:

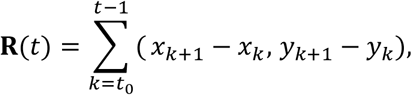

and defined the predicted home vector as 𝐇(𝑡) = −𝐑(𝑡). For each frame during the return phase, we computed the angular difference between the instantaneous movement direction and the home vector direction. A home-alignment index (HAI) was calculated for each bout as the circular concentration of these angular differences, with values approaching 1 indicating strong alignment to the home vector and values near 0 indicating random orientation.

To control for non-PI explanations, we generated heading-shuffle controls by permuting heading increments within each bout while preserving step lengths. HAI values from real data could be compared to shuffled distributions to unmask a nonrandom pattern. We averaged bout-level PI metrics (return error ∣ Δ𝑠 ∣and HAI) within fish for early (Minute 1) and later (Minute 8) exploration. Comparisons were performed within fish using paired, non-parametric statistics (Wilcoxon signed-rank tests or paired permutation tests where appropriate).

#### 2.4.5. Return-to-anchor spatial specificity

To quantify spatial specificity of returns, we computed the boundary coordinate 𝑠_return_ at the first boundary re-contact following the outbound phase. We defined “return error” as the wrapped difference Δ𝑠 = 𝑠_return_ − 𝑠_0_. For each fish, the distribution of Δ𝑠 across bouts was compared to a within-fish shuffle control in which anchor labels were randomly permuted across bouts (10,000 permutations). Spatial clustering was quantified using the mean resultant length of the circular error distribution, and significance was assessed by comparing observed clustering to the shuffled null distribution.

#### 2.4.6. Corrective-turn model from signed body-axis heading

We used body-axis orientation to compute signed heading updates Δ𝜃_𝑡_ = wrap(𝜃_𝑡+1_ − 𝜃_𝑡_) for each fish and minute. For each sharp-turn event, we identified the preceding 2–3 wall touch onsets within a 150-frame window (same minute) and required at least one Reorient frame between the final WT onset and the SharpTurn. We defined the accumulated heading change Θ_accum_ = ∑Δ𝜃 from the first qualifying WT onset up to (but not including) the SharpTurn frame. The predicted corrective turn was Θ_pred_ = −(180^∘^ − Θ_accum_) (wrapped to [−180^∘^, 180^∘^]), and the observed corrective turn was the signed Δ𝜃 at the SharpTurn frame. Residuals were computed as wrap(Θ_obs_ − Θ_pred_) and summarized using circular mean and circular standard deviation.

## 3. Results

### 3.1. Swim locations and dynamics

If fish engage in a systematic behavioral strategy to learn about their immediate space, we should expect to see a nonrandom pattern of swim locations with the novel tank and a change over time in the expression of any postural and swim dynamics as they familiarize themselves to their new space (Fig. S3). This may arise from their tank boundary-related interactions and/or a shift in the postural and swimming characteristics. We used the x-y coordinate data to track turning behavior from heading transitions, wall contact events (see Methods) and swim-locations, such as mean distance from the tank boundary. The relative distribution of the various body postures and actions (swim/glide) states over time are shown as pie charts in figure S4. These data show that on average the overall proportion of time fish spend waiting decreases and the swim increases. The occurrence of reorient events also increases with time.

A comparison of the distributions of distances between the OC marker on the head and the tank boundary showed an increase in the relative (to each time segment) frequency of locations where a fish was present near the boundary later in time (Fig. 1A). A proportion of densities plot indicated that across time zones on average, fish spent slightly less time near the boundary in the first minute compared to later (minutes 5 and 8) and accordingly a higher proportion of presence near the center is noted in the first minute (Fig. 1B). One reason for this may be that fish were released near the center of the tank where they lingered a bit before swimming towards the tank boundary. The space exhibited roughly 1 cm long wavelength of peaks and valleys of occupation over the first and fourth minute, suggesting preference for regional bands for either stationary presence or swim activity. This plot does not tell us, however, whether there were multiple excursions to the boundary in one time segment vs. others and/or whether they were stationarily positioned or swimming around the periphery. To clarify such differences, if any, we examined the WT event rates over time (Fig. 1C) and changes in turn angle close to the boundary vs. when swimming within the mid-region of the tank (Fig. 1D). The wall touch events remained relatively stable, oscillating slightly over two-minute timeframes. The turning behavior, however, was quite different during their presence near vs. away from the tank wall (Fig. 1D). The data show a significant (P < 0.001) early vs. late decline in turn angles near the wall and a significant (P < 0.001) rise in turn angles over time when swimming away from the wall. A large spread of variation in turn angles explain the low correlation coefficients.

### 3.2. Postural and swim pattern analysis

To standardize postural data definitions, we list component postures and actions and indicate differences in the parameters defining each posture, such as reorienting vs shsrrp turn (Table 1). We only use this as an *a priori* qualitative description of postures and action patterns and not to directly test for differences over time.

**Table 1.**
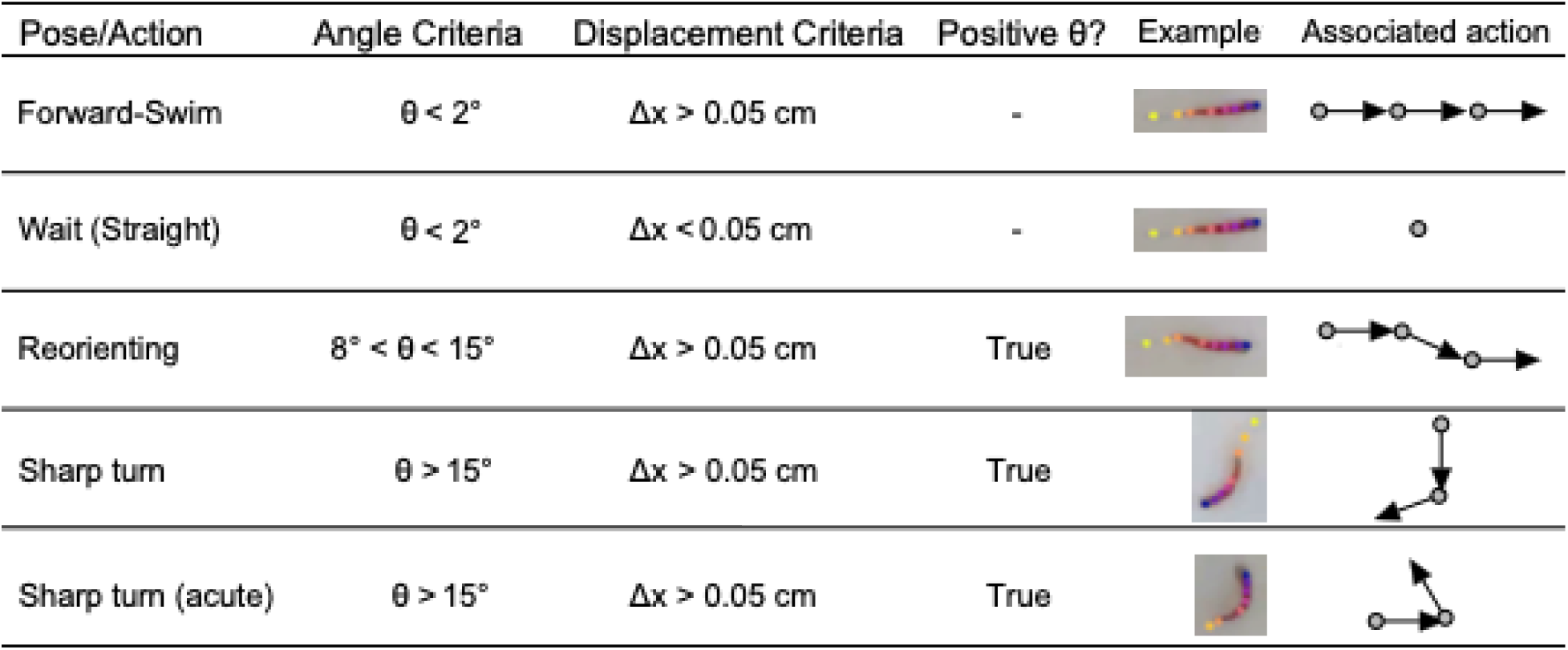
Postural classification. List of the given names of pses and related actions identified by B-SoiD after introducing an animal into the novel tank environment. The column on the right shows a cutout of the screenshot of each pose in an adult zebrafish. We also list three criteria used in the pose analysis. Body angles were measured using markers at the ocular center (OC), the mid-flank (center-point) and tailfin-base. Body angles between 8 and 15 deg. were labeled as “slight turns”.

| Pose/Action | Angle Criteria | Displacement Criteria | Positive $\theta$? | Example | Associated action |
| --- | --- | --- | --- | --- | --- |
| Forward-Swim | $\theta < 2^\circ$ | $\Delta x > 0.05$ cm | - | | |
| Wait (Straight) | $\theta < 2^\circ$ | $\Delta x < 0.05$ cm | - | | |
| Reorienting | $8^\circ < \theta < 15^\circ$ | $\Delta x > 0.05$ cm | True | | |
| Sharp turn | $\theta > 15^\circ$ | $\Delta x > 0.05$ cm | True | | |
| Sharp turn (acute) | $\theta > 15^\circ$ | $\Delta x > 0.05$ cm | True | | |

### 3.3. Behavioral state classification and event encoding

To quantify how exploratory strategies reorganize with experience, we modeled behavioral dynamics as state-transition networks for the early (first) and late (eighth) minutes of exploration across seven individuals. In two individuals, recordings were terminated after 5 minutes since they did not show significant activity. States were defined after thresholding for different ranges of quantitative data on swim and turn parameters and calculated transitions from the frequency with which one state changed to another.

To compute a sharp turn, we combined conditional postural state with thresholding of continuous values for head transition angles. Swimming behavior was quantified from frame-wise tracking data and discretized into mutually exclusive postural–kinematic states: Glide, Wait, Swim, Reorient, SharpTurn, and AboutTurn. State assignment followed a hierarchical precedence rule: AboutTurn (rare) → SharpTurn → Reorient → Swim → Glide → Wait (common), ensuring that high-curvature turning events were not masked by concurrent and common locomotor activity. WT was treated as a *state modifier* rather than a primary behavioral state. WT events were defined using a conservative geometric criterion based on proximity to the rounded tank boundary (distance to tank wall ≤ 4 mm). WT could co-occur with any behavioral state and was not permitted to override state identity.

### 3.4. Early to late-phase state transitions

To illustrate experience-dependent changes in behavioral transition structure, we plot Markov networks for the first minute and a difference network between the eighth and first minute of activity (Figs. 2A and 2B). In the first minute, the network is dominated by frequent transitions between wall touch, reorient, and high-curvature turn states, with thick edges linking wall contact to reorientation and sharp or about turns, consistent with boundary-anchored exploratory sampling (Fig. 2A). By minute 8, overall network structure is preserved but edge weights are redistributed, with reduced dependence of turn states on wall contact and increased transitions among swimming and turning states away from the boundary. These changes are made explicit in the Δ network (minute 8 − minute 1). This allowed us to directly test: if a turn occurred, was it wall-associated? Notably, transitions from WT to Reorient and from WT to sharp turns, including about-turns, are selectively reduced, whereas transitions among turn and locomotor states independent of wall contact are enhanced. In short, Markov analysis revealed structured reorganization of behavioral dynamics over time with the difference network demonstrating selective strengthening and weakening of specific transitions rather than uniform scaling across all states.

**Figure 2.**
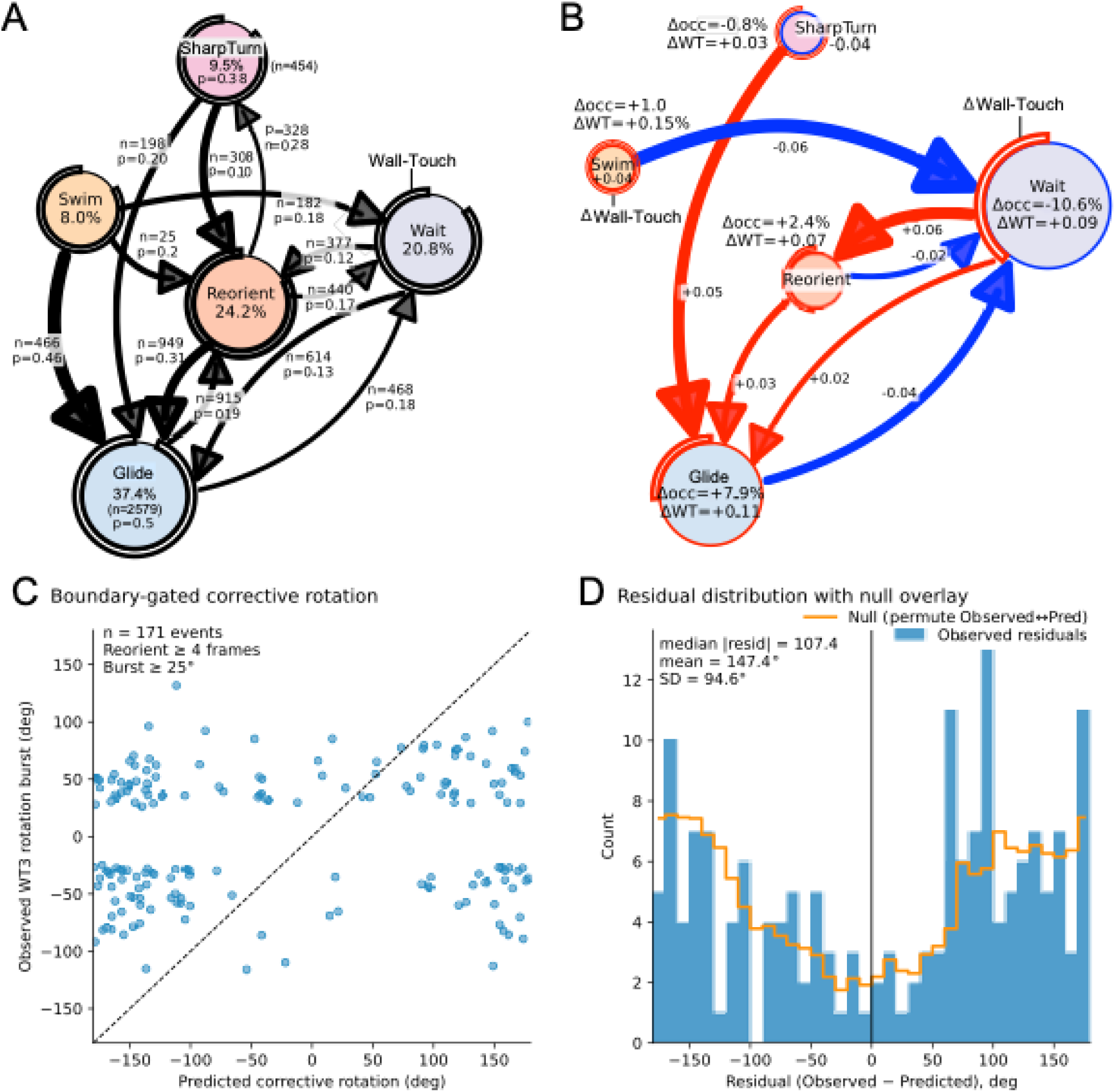
A. Markov transition network during Minute 1 of exploration reveals strong coupling between boundary contact and turning-related states, with high-probability transitions linking Swim, Reorient, and SharpTurn. Outer arc lengths indicate substantial overlap between Reorient/SharpTurn states and WT, consistent with boundary-triggered orientation updates. B. Difference network (Δ[8–1]) shows a redistribution of transition probabilities over time, with reduced reliance on WT-associated turning and increased transitions among non-boundary states. C. Boundary-anchored corrective rotation links state-space dynamics to event-level path integration. D. Event-level analysis demonstrates that discrete heading updates at boundary contacts and a dominant reorientation bout predict a corrective rotation occurring during the third wall contact. Each point represents a wall-touch (WT3)-gated corrective event across fish. Orange line represents the difference Markov network (Minute 8 − Minute 1) showing a reduction in transitions coupling WT to Reorient states, despite persistence of turn states later in exploration. See Figure S5 for a conceptual schematic summarizing the proposed mechanism: repeated boundary contacts accumulate angular error through discrete heading updates and reorientation, which is compensated by a corrective turn at the third wall encounter.

Difference networks highlighted increased stability within locomotor states and altered coupling between turn states and sustained swimming, indicating experience-dependent refinement of action sequences (Fig. 2B). Reciprocal transitions between turning and locomotor states were asymmetric, suggesting directional biases in behavioral progression rather than reversible switching. WT overlap decreased selectively for high-curvature turn states over time, consistent with a progressive reduction in boundary-driven reorientation. To visualize the coupling between turn states and wall contact, turn nodes are annotated with colored arcs (concentric circular lines) whose lengths indicate the fraction of each state overlapping with WT. Arc lengths are longest for reorientation and sharp turns early in exploration but are markedly reduced by minute 8, indicating a decoupling of directional reorientation from boundary contact over time. Together, these results reveal a systematic reorganization of exploratory state dynamics in which early wall-anchored turning gives way to more internally structured transitions, consistent with the emergence of path-integration-like control of movement. Thus, consistent with the Δ(8–1) Markov network, which shows reduced transition coupling between turning states and boundary-associated contexts, WT-conditioned analyses reveal that SharpTurn events become significantly less associated with wall contact over time, despite preserved turning state occupancy (n = 7,129 WT→Reorient→SharpTurn sequences (2–3 WT onsets, within 150 frames, same minute).

### 3.5. Reorganization of behavior

To assess whether exploratory behavior reorganizes over time, we compared behavioral state occupancies and transition probabilities between Minute 1 and Minute 8 using paired within-fish statistical tests. Several transitions central to the hypothesized corrective turning sequence exhibited significant changes across fish (Fig. 1B). In particular, the probability of the Reorient → SharpTurn transition was significantly reduced between early and late exploration (Δ = −0.069, Wilcoxon signed-rank test, *W* = 1, *p* = 0.016; Table S1), corresponding to a prominent blue edge in the Δ(8–1) Markov network (Figure X). Similarly, the Swim → Reorient transition showed a significant decrease over time (Δ = −0.055, *W* = 2, *p* = 0.031; Table S1).

At the state level, turning behaviors became progressively less coupled to boundary contact. The fraction of Reorient frames overlapping WT was significantly reduced from Minute 1 to Minute 8 (Δ = −0.242, *W* = 0, *p* = 0.008; Table S1), with a similar trend observed for SharpTurn (Table S1). These changes indicate a decoupling of reorientation and turning from immediate tactile cues as exploration proceeds. Together, these paired statistical results validate the transition reweighting visualized in the Δ(8–1) Markov network and demonstrate that exploratory behavior undergoes a structured reorganization from boundary-anchored turning toward increasingly internalized control of behavioral state transitions. This reorganization parallels the emergence of corrective sharp turns that compensate for accumulated reorientation error, suggesting that early boundary-driven updates give rise to internally coordinated navigational control.

#### 3.5.1. Experience-dependent changes in average fish-normalized behavioral composition

Other computed parameters include Mean wrapped residual: −173.1°, Circular SD: 60.7°, and Median |residual|: 143.1°. Across 7,129 WT→Reorient→SharpTurn sequences, signed body-axis heading revealed structured corrective-turn dynamics following clustered boundary contacts (Fig. 2C). Specifically, the magnitude and direction of SharpTurn-associated heading changes were systematically related to the accumulated heading change preceding the turn, consistent with a discrete correction step operating after boundary-gated reorientation.

However, residuals remained offset from zero under this initial window definition, suggesting that the correction rule is implemented over a more restricted subset of angular updates (e.g., WT- and/or Reorient-specific increments) and/or that SharpTurn rotations extend over multiple frames beyond the event marker. In short, consistent with the Δ(8–1) Markov network, which shows reduced transition coupling between turning states and boundary-associated contexts, WT-conditioned analyses reveal that SharpTurn events become significantly less associated with wall contact over time, despite preserved turning state occupancy.

#### 3.5.2. Experience reduces boundary coupling of turning behavior

Paired comparisons were performed to assess the probability that a turning event co-occurs with wall contact, (WT ∣ Turn), for Reorient and SharpTurn states in early (Minute 1) and later (Minute 8) exploration (Fig. 2D). Each dot represents one fish; gray lines connect paired observations across minutes. Large symbols indicate the mean ± SEM across fish. Statistical comparisons were performed within fish using two-sided Wilcoxon signed-rank tests; exact p-values are annotated above each category. For Reorient, all paired differences were zero (all fish had 𝑃 (WT ∣ Reorient) = 1 at both time points), precluding statistical testing. Together, the data show that turning behavior persists across time but becomes less tightly coupled to boundary contact, particularly for SharpTurn events. Together, these analyses indicate a transition from boundary-anchored navigation early in exploration to boundary-independent turning strategies later, consistent with a path-integration–like process.

#### 3.5.3. Path integration-like navigation dynamics

Here we asked the question, do fish return to the same boundary location they last touched more than expected by chance? We used body-axis orientation to quantify signed angular updates, independent of translational noise (Fig. 3A). This allowed us to conclude that reorient events accumulate angular error, SharpTurns act as discrete corrective operations, correction magnitude is predicted by integrated heading error, and boundary contact gates when this correction is triggered (Fig. 3B). In short, we can conclude that path integration is not just an observed behavioral phenomenology but occurs at the algorithmic level in adult zebrafish. Figure 3C shows the fraction of Reorient and SharpTurn events that co-occur with WT in early (Minute 1) and late (Minute 8) exploration. Each dot represents one fish; lines indicate paired comparisons. A downward shift in paired dots immediately conveys “decoupling. Although turning states persist across time, their association with wall contact is significantly reduced with experience, indicating decoupling of corrective turning from boundary interaction.

**Figure 3.**
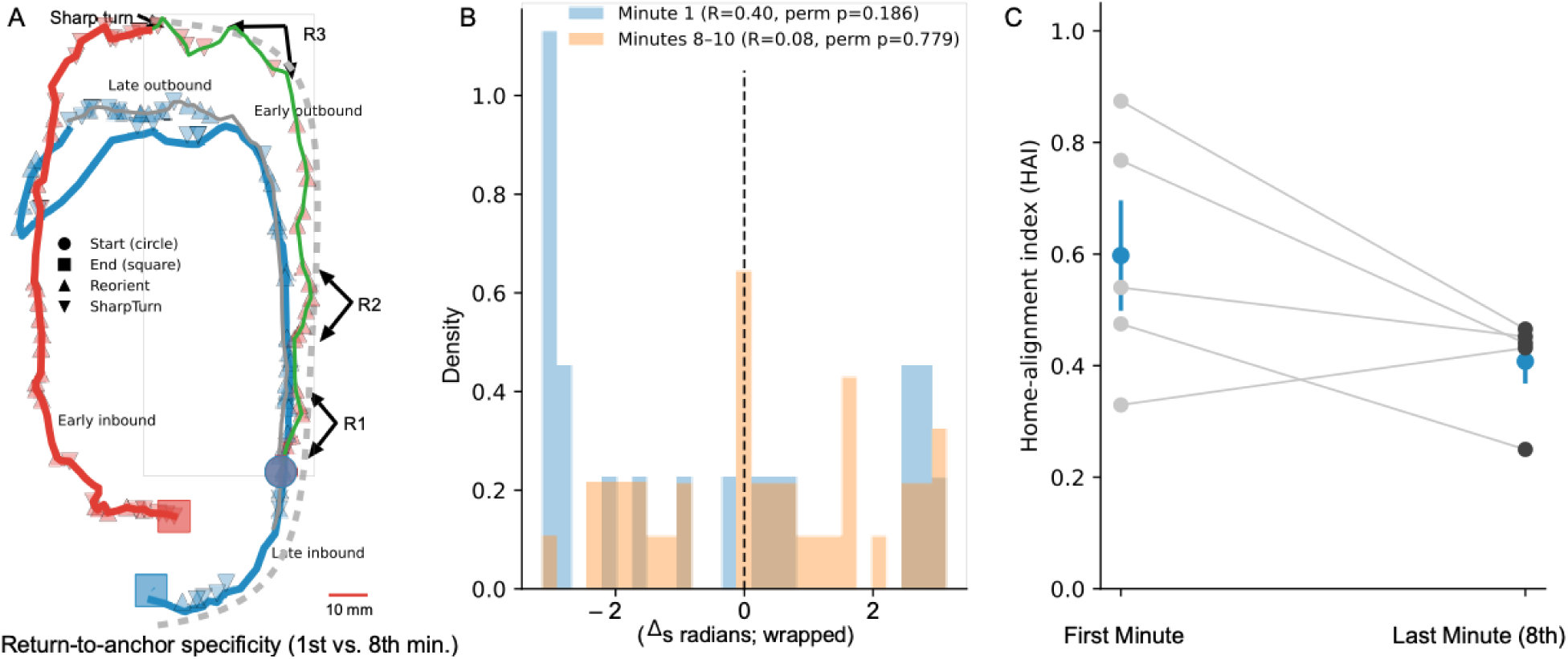
Wall-contact–anchored bouts reveal PI-like navigation in adult zebrafish (A) Example excursions anchored at WT events. Outbound (thin) and inbound (thick) trajectories are shown with predicted home vectors computed by integrating frame-wise displacement from the anchor. (B) Population distribution of return-location error along the boundary (Δ𝑠) relative to the anchor (0), compared with a within-fish shuffle control. (C) Experience-dependent improvement in PI-like performance quantified by return-location error (or home-alignment index) from Minute 1 to Minute 8/10, paired within fish.

For a definitive confirmation that experience reduces boundary coupling of turning behavior, we made paired comparisons of the probability that a turning event co-occurs with wall contact, 𝑃(WT ∣ Turn), for Reorient and SharpTurn states in early (Minute 1) and later (Minute 8) exploration. Each dot represents one fish; gray lines connect paired observations across minutes. Blue symbols indicate the mean ± SEM across fish. Statistical comparisons were performed within fish using two-sided Wilcoxon signed-rank tests; exact p-values are annotated above each category. For Reorient, all paired differences were zero (all fish had 𝑃 (WT ∣ Reorient) = 1 at both time points), precluding statistical testing. Together, the data show that turning behavior persists across time but becomes less tightly coupled to boundary contact, particularly for reorientations followed by SharpTurn events.

### 3.6. A Swim-Touch-Reorient (STR) sequence

When fish are first introduced into the novel environment, they sometimes swim alongside the edge of the tank, but typically they exhibit a consistent swim pattern or maneuver: swimming forward until they contact the edge. This is followed by a re-orientation or sharp turn ending in a second contact and subsequently a third contact with the edge (labelled as touchpoints). All fish repeatedly engaged in this common or stereotypic maneuver during the early period of tank exploration. We summarize the common characteristics of this maneuver initiated by swimming forward as a bout of swim-touch-reorient (STR) sequences. The last contact within each STR bout is followed by a near-about turn (classified as a “sharp turn” considering a nondirection body-axis orientation) away from the tank wall to return to the neighborhood of the initial (anchor) WT point, after which this entire action-sequence is repeated from a new start location, typically close to previous one. Figure 4A shows an example of a STR sequence with the track traced within a bubble plot of fish locations (x-y coordinates). Per the statistical analysis and visual observations, STR bouts declined or were absent during later minutes of recording as illustrated in figure 4B.

**Figure 4.**
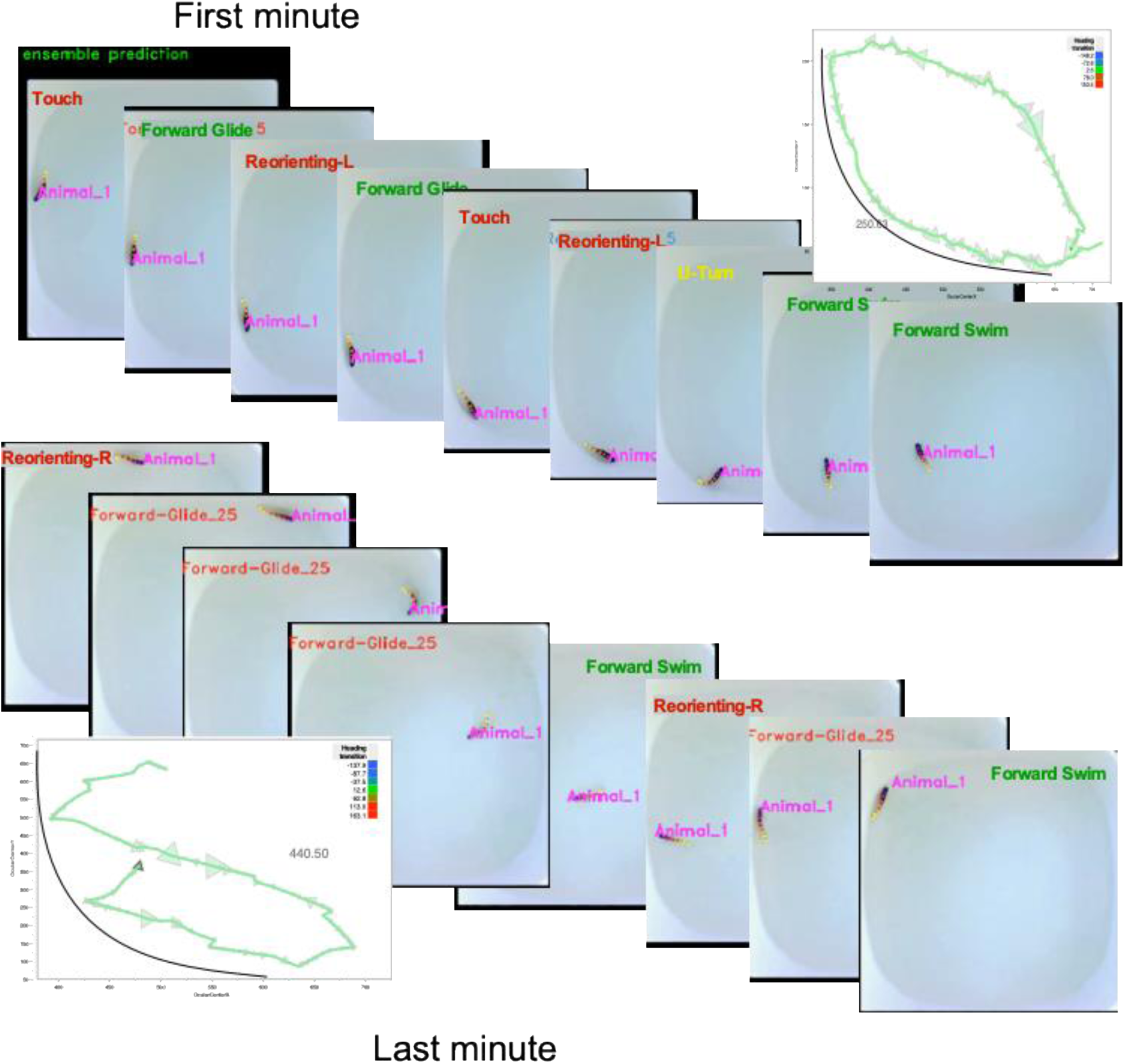
Stacks of video-fames and bubble plots (top-right and bottom-left corners) illustrating behavioral strategy adopted pre- and post-mapping of behavioral space. Initially, two or more successive contacts with tank-wall were followed by return to the initial location (top panel). Later, fish tended to without reorient-coupled wall contacts (bottom panel).

### 3.7. Development of the STR sequence

We were curious if zebrafish engage in swim bouts containing STR sequences during the embryonic stages and if not, when does this behavior first develop. For this, we tracked development of the stereotypic STR sequences throughout larval development, from 7 to 44 days post-fertilization (dpf). We observed that across developmental stages, forward-glide and reorienting movements make up the majority of movement bouts; however, wall touch frequency increases as embryonic development progresses (Figure 7A). A go swim-and-touch bout is essential for providing sensory touch information for PI strategy. At 30-44dpf, we see a significant increase in wall touching behavior (student’s t-test; p = 0.0122) compared to 7-13 dpf (Figure 5A), while other bout types show no significant variation (ANOVA; df = 3, p > 0.05) across developmental stages. We further show a marked shift in the transition matrix of movement bouts as zebrafish progress through development (PERMANOVA; Bray-Curtis, *R^2^*= 0.48, *df* = 3, *p*<0.05).

**Figure 5.**
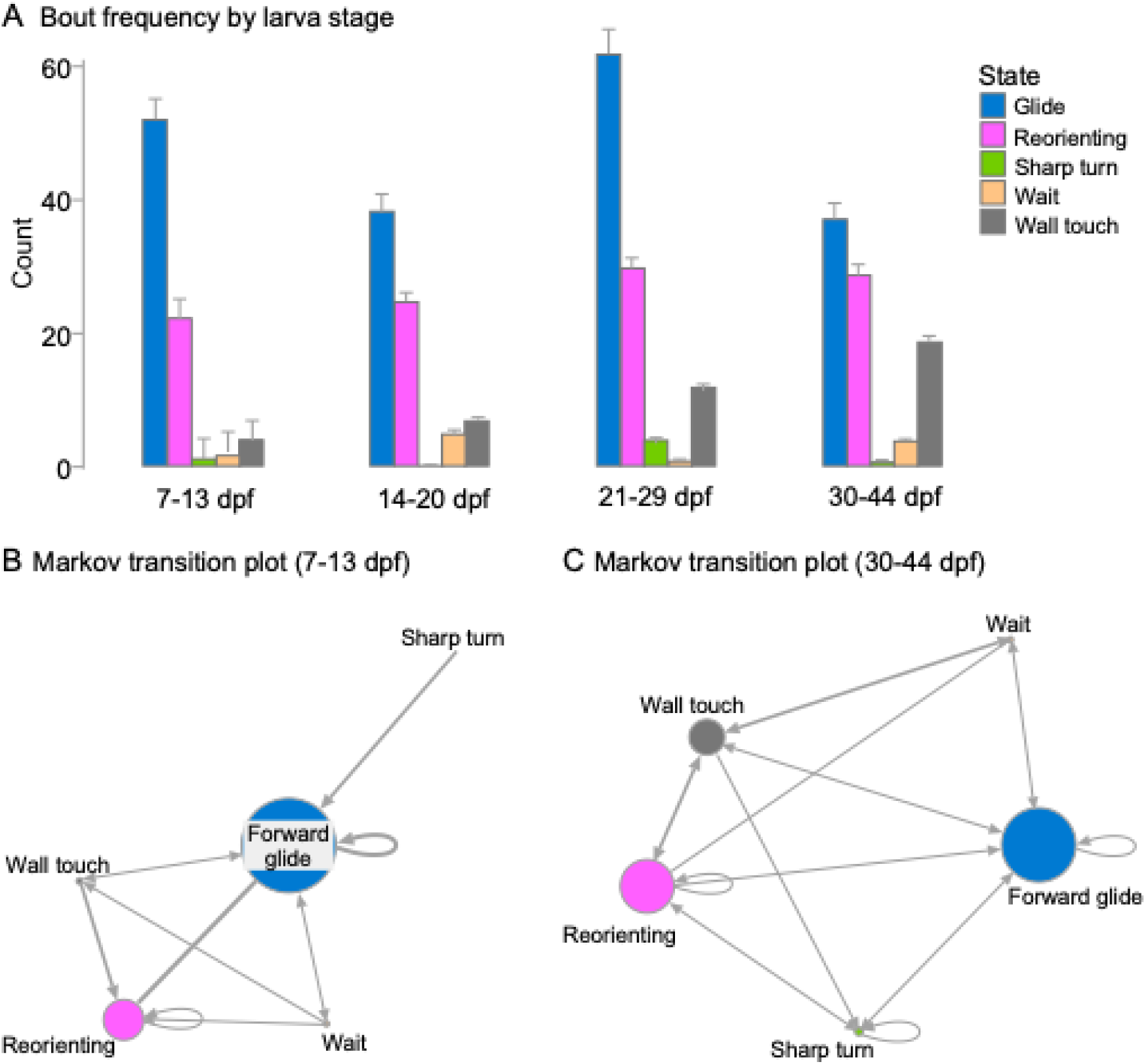
Exploratory swim behavior differs by developmental stage during the first minute of swimming. (A) Bar graph of frequency of each kind of movement bout, by developmental stage days post-fertilization (dpf). (B) Markov transition plot of larval zebrafish 7-13 days post-fertilization. (C) Markov transition plot 30-44 days post-fertilization. Node size corresponds to frequency of movement bout. Thicker arrows denote greater frequency of transitions. Plots were made from 60 seconds of observation of (n=9) zebrafish larvae (6 fish at 7-13dpf; 3 fish at 30-44dpf), normalized across fish. dpf: days post-fertilization.

Pairwise comparisons between fish 7-13 dpf and 30 – 44 dpf showed significantly different bout transition matrices (PERMANOVA, Bray-Curtis, *R^2^* = 0.13, *df* = 1, *p*<0.001). At 7-13 dpf, fish primarily execute transitions from forward glide to reorienting, or forward glide to a repeat forward glide, as they navigate the novel environment confined to bout types (Shannon entropy, *H* = 2.46) without a distinct strategy. There is little involvement in dedicated sharp turn movements or exploratory touch behavior on the tank boundary (Figure 5B). Whereas in fish at four or more weeks of age post-fertilization, we find that transitions are less concentrated in recurring forward glide bouts, with significantly fewer transitions from forward glide to forward glide (Welch’s t-test, *p*=0.008) and forward glide to reorient motifs (Welch’s t-test, *p* = 0.045) compared to 7-13 dpf. Bout transitions are instead more evenly interspersed (Shannon entropy, *H* = 3.50) *between* touch, reorienting, forward glide, and sharp turn movements, consistent with the development of PI-like STR maneuvers (Figure5). This critical behavior for spatial navigation is innately developed from larval stages – an evolutionary solution for self-orientation and survival.

## 4. Discussion

In this study, we observed repeated patterns of activity with a bias towards swimming near the tank boundary in contrast to random sampling of locations in the same environment. This type of anxiety-driven behavior is commonly observed in rodents and other species (Seibenhener and Wooten, 2015). A closer examination revealed a systematic exploration of the novel environment. We compared swim patterns during the early and late phases of swimming over a 8 to 10-minute period after fish were placed in the tank. Given that this study were conducted in ambient light, albeit visually cueless environment, there is a slight possibility of the fish cueing in on some type of light distribution cue to gain familarity with their surroundings. Ongoing studies in complete darkness using infrared videography, however, show that zebrafish exhibit similar behaviors while exploring a novel tank environment (Shraybman and Kanwal, 2024).

### 4.1. A spatial learning strategy - boundary-anchored exploration scaffolds path integration–like navigation

We discovered that when zebrafish are placed in either a visually uninformed or deprived environment, they immediately engage in a systematic exploration behavior. Our results indicate that wall-contact behavior in zebrafish is not adequately described by a single scalar measure such as contact frequency or dwell time. Instead, boundary interactions occupy a low-dimensional strategy space, with individual animals adopting distinct and stable modes of interaction that evolve over time. The absence of a uniform population-level shift, together with pronounced and coherent within-animal trajectories, suggests that boundary behavior reflects individual-specific strategies rather than a common temporal progression. Some fish consistently revisit a preferred boundary location, while others progressively patrol the tank perimeter, and these tendencies persist even as overall engagement with the wall changes. This observation aligns with emerging work across species discussed later showing that exploratory and navigational behaviors are best understood as structured phenotypes rather than noisy realizations of a shared mean. By unwrapping spatial trajectories and applying circular metrics, our approach makes these phenotypes explicit and quantifiable. In this framework, behavioral change is naturally represented as movement through strategy spac**e**, rather than as an increase or decrease in a single variable.This perspective opens several avenues: linking boundary strategies to neural circuit dynamics, internal state, or learning; interpreting patrolling versus focal revisitation as distinct navigational modes; and extending the strategy-space framework to other spatial behaviors and species. More broadly, these results argue for analytical approaches that preserve individual structure when studying complex behavior.

Our analyses reveal that exploratory behavior in adult zebrafish is not static but undergoes a statistically robust reorganization over time. Early in exploration, turning-related states are tightly coupled to boundary contact, as evidenced by high WT overlap and strong transition probabilities linking Swim, Reorient, and SharpTurn states. Paired within-fish statistical testing confirms that these boundary-anchored transitions—most notably the Reorient→SharpTurn and Swim→Reorient transitions—are significantly reduced as exploration proceeds (Table S1). This reweighting of transition probabilities is reflected directly in the Δ(8–1) Markov network, in which turning-related edges decrease while transitions among non-boundary locomotor states increase. Importantly, this reorganization cannot be explained by a simple reduction in activity or wall-following. Instead, the decrease in WT overlap for Reorient and SharpTurn states, together with preserved turning structure, indicates a decoupling of orientation updates from immediate tactile cues. Mechanistic analyses further show that early sequences of repeated reorientation following boundary contact culminate in corrective sharp turns that compensate for accumulated angular error, allowing fish to return toward their initial contact location. Over time, these same turning states occur increasingly without boundary contact, suggesting that boundary-derived information becomes internalized and used to guide navigation in the absence of direct sensory input.

These results support a model in which boundary contact initially scaffolds spatial updating, providing external reference points that anchor self-motion cues. As exploration progresses, zebrafish increasingly rely on internally coordinated state transitions to guide movement, consistent with a path integration–like strategy. While we do not claim the formation of a full metric cognitive map, the observed behavioral reorganization demonstrates that adult zebrafish integrate idiothetic and tactile information to estimate spatial relationships in a novel environment, positioning this model system as a powerful vertebrate platform for dissecting the neural and genetic basis of navigation.

Together, our analyses indicate that experience reorganizes behavior by stabilizing state persistence and reducing boundary-driven corrective transitions, rather than suppressing exploration outright. Markov structure reveals changes in *how* behaviors are sequenced, while event-based averages reveal changes in *how often* transitions are initiated. The convergence of these measures supports a model in which spatial experience promotes smoother, more predictive locomotor strategies with reduced reliance on high-curvature reorientation. Briefly, within a matter of a few minutes, the transition architecture among locomotor and turning states is reorganized in a manner consistent with experience-dependent refinement of spatial behavior. Early exploration (Minute 1) exhibited higher coupling between locomotion and high-curvature reorientation, together with elevated WT overlap, indicative of frequent boundary-driven corrective maneuvers when the internal estimate of position or heading is still unstable. By minutes 8–10, the Markov structure showed a shift toward increased persistence within locomotor states and selective attenuation of turn-linked transitions, while WT became less associated with turning states. This pattern is consistent with improved path integration–like stability: as animals accumulate self-motion evidence and learn the geometry of the arena, the mapping between idiothetic cues (e.g., swim bouts and turns) and expected sensory consequences becomes better calibrated, reducing the need for abrupt reorientation and boundary contacts to resolve accumulating error. Importantly, the divergence between duration-based occupancy and minima-based initiation metrics suggests that experience does not simply suppress exploration; rather, it consolidates motor sequences into longer, smoother bouts with fewer discrete corrective turn initiations—an expected signature of a system transitioning from reactive boundary control toward predictive, internally guided navigation.

### 4.2. Path Integration across species

Path integration has been studied in a few but wide variety of species, including humans (Aharon et al., 2017; Bostelmann et al., 2020; Etienne et al., 1986; Heinze et al., 2018; Mittelstaedt and Mittelstaedt, 1980; Poll et al., 2016; Stone et al., 2017; Wiltschko and Wiltschko, 1978). PI was first observed and experimentally tested in desert ants (Lebhardt and Ronacher, 2014; Müller and Wehner, 1988; Wehner and Srinivasan, 1981; Wittlinger et al., 2006). Since then, map-like spatial memories and mechanisms for heading and orientation have been elucidated in honeybees, and more recently in fruit flies, to understand the neural basis of orientation and PI (Behbahani et al., 2021; Heinze et al., 2018; Hulse et al., 2021; Menzel et al., 2005). Honeybees can even communicate distance and direction information to conspecifics through dance-like movements that have been studied in great detail since their discovery (Ai and Farina, 2023; Menzel et al., 2005; Riley et al., 2005). PI and its neural basis has been studied in great detail in rats and more recently in humans (Alyan and McNaughton, 1999; de Cothi et al., 2022; Langston et al., 2010; Moser et al., 2008). Studying PI in a non-mammalian vertebrate model species can advance our understanding of the neural origins of spatial navigation and eventually of its neurogenetic basis.

PI represents a key mechanism by which animals maintain an internal representation of their spatial location relative to starting point to compute a home vector (Conklin and Eliasmith, 2005; Vickerstaff and Di Paolo, 2005). The main function of PI is to allow an organism to compute their current position by integrating directional and distance cues obtained during movement. Ideally, this navigational strategy alone can enable an animal to return to their point of origin without relying on external landmarks (Etienne and Jeffery, 2004). Homing pigeons continuously update their location by integrating movement-related information and initially rely on PI before incorporating external cues to refine their navigation (Bingman and Able, 2002; Holland, 2003). For other areal species, such as bats, visual information is severely restricted. They can return to a starting point after complex foraging routes even in the absence of external cues (Geva-Sagiv et al., 2015). As with the bat study, in our experimental setup, animals introduced into a novel tank environment had no clearly demarcated reference points along the boundary of the test tank other than a static overhead camera face that, given its size and short distance from the water surface, did not serve as a useful marker. Recent studies of fish navigation suggest that, like terrestrial species, fish, including zebrafish, use self-motion cues to construct a map of their surroundings and compute return paths, particularly in visually limited conditions (Alexander et al., 2022; Rodríguez et al., 2021). Cichlid fish (*Pelvicachromis pulcher*), maintain an internal record of movement direction and distance to return to a home shelter (Sibeaux et al., 2024).

PI was first demonstrated and extensively studied in Saharan desert ants (*Cataglyphis*) as a model of efficient navigation in featureless environments (Wehner and Srinivasan, 1981). These ants can monitor the distance traveled by counting their steps (Wittlinger et al., 2006). When physically displaced, desert ants continue to walk in the direction of their nest’s expected location rather than adjusting to a new one, underscoring the importance of internal cues for PI (Collett and Collett, 2000). To determine direction in their natural environment, however, they may build tall nest hills and use celestial cues such as the sun’s position and polarized light patterns from constellations in the sky (Freire et al., 2023) Winner, 2003). Using this combination of cues, they can compute the return path to their nests after traversing vast and barren desert landscapes (Müller and Wehner, 1988). Similar strategies have been observed in honeybees (*Apis mellifera*), which must efficiently navigate between their hive and foraging locations. Bees update their position relative to the hive by continuously tracking their flight path, a process facilitated by optic flow (Esch and Burns, 1995; Srinivasan et al., 1996). This internal measure of distance, coupled with directional information from the sun, enables precise navigation between food sources and the hive (Esch and Burns, 1995). PI has also been demonstrated in *Drosophila* where it depends on a combination of visual and olfactory cues (Strauss, 2002). In a relatively small, contained environment, local visual landmarks and boundaries become important and must be combined with PI to create neural maps of one’s immediate environment. We propose that the purpose of STR maneuvers, observed in our study, is to combine external and internal cues with PI to map the geometry of the tank environment. Once this is accomplished, the use of STR maneuvers becomes irrelevant.

### 4.3. Navigation within three-dimensional space

Most studies on PI are focused on animal movements in a largely two-dimensional environment. Studies on aerial species, such as birds and bats, offer a unique perspective on how PI may allow navigation in three-dimensional space (Kothari et al., 2018). Studies show that bats integrate self-motion cues with a combination of sensory inputs, including echolocation and proprioception, to maintain an internal representation of their position during flight and perform PI (Aharon et al., 2017; Wohlgemuth et al., 2016). Aquatic, like aerial environments, are inherently three-dimensional and this adds another layer of complexity in the natural environment. Regardless, for some birds, like eagles that soar in the sky, a two-dimensional view of visual landmarks may be sufficient to map their operational space. Although fish too need to map their space within three dimensions, we focused on a two-dimensional plane in this study. Because of the relatively shallow water surface in our setup, the depth dimension was not relevant as we needed to keep them within the depth-of-field of focus of the camera and allow markerless tracking using DLC. For zebrafish that typically thrive in shallow waters, two-dimensional PI may be sufficient to map their environment. The depth dimension may be independently tracked visually within a two-dimensional frame-of-reference since zebrafish have good color vision, and the same visual cues are largely visible from a short range of depths (Baden, 2021; Fornetto et al., 2020; Sato and Kefalov, 2025).

### 4.4. Spatial mapping in zebrafish: from “see-and-go” to a “go-and-touch” strategy

As with well-studied mammalian species, in the natural environment, zebrafish can associate exterosensory (visual) cues with interoceptive (proprioceptive, temporal and vestibular) information about turns and swim distances to create a neural representation of their space (Angelaki et al., 2011; Campos et al., 2014; Dunn et al., 2016; Kaski et al., 2016). In the presence of visual cues, this can be labeled as a “see-and-go” strategy for mapping space (Fig. 6A). In the absence of useful visual cues, zebrafish still have access to swim-generated idiothetic cues, such as proprioception, water flow over its body surface as well as recurrent feedback from motor activity triggering tail undulations. Based on our classification and analysis of observed fish behavior patterns, we propose that in the absence of a clearly visible boundary or visual landmarks, zebrafish use a systematic “go-and-touch” strategy to map their home space (Fig. 6B). A key feature of this map-making strategy involves a swim forward to contact the tank wall followed by a reorientation of head direction. A series of these STR maneuvers (typically three) followed by a short path back to the first wall contact point demonstrates the ability of zebrafish to accomplish PI. This behavioral repertoire represents a reversal of the sensorimotor sequence into a motor-sensory stop sequence where the touch may trigger a time-tracked, motor/proprioceptive memory consolidation event.

**Figure 6.**
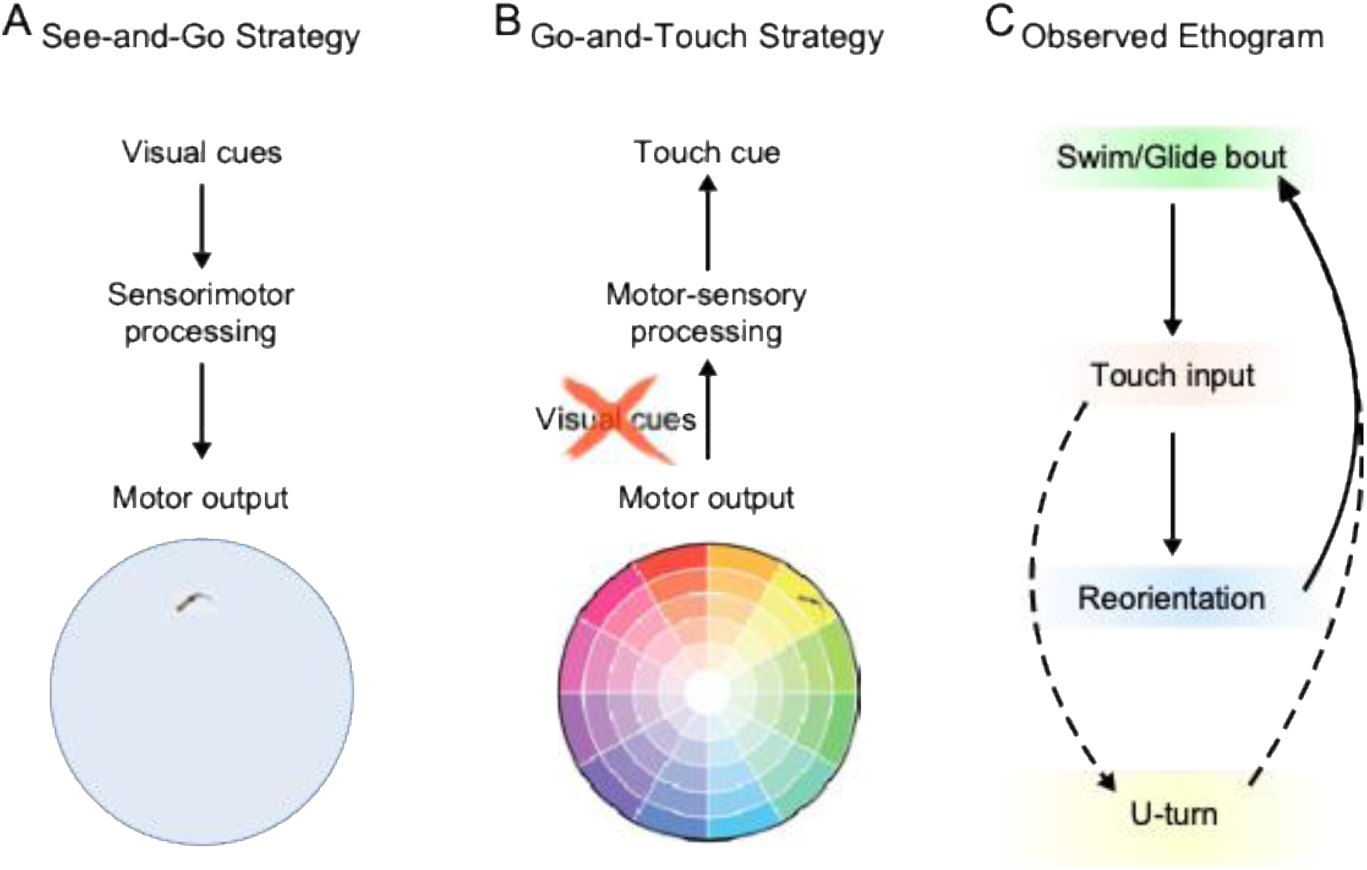
Flow chart of behavioral strategy for path integration. In their natural environment, zebrafish match idiothetic information with allothetic cues generated via optic flow and other mechanisms to generate a cognitive map. B. In the absence of visual information, zebrafish swim along a straight path until they make contact. The touch input triggers a minor reorientation to swim away from the boundary until a new contact point is encountered. Repeated sequences of swim-touch-reorientation bouts continue until a cognitive map, denoted by a color wheel, is created within which the fish can locates its position. C. Ethogram of the STR maneuver (solid lines) and PI behavior (dashed lines) with arrows pointing to the two components of the map generation paradigm.

**Figure 7.**
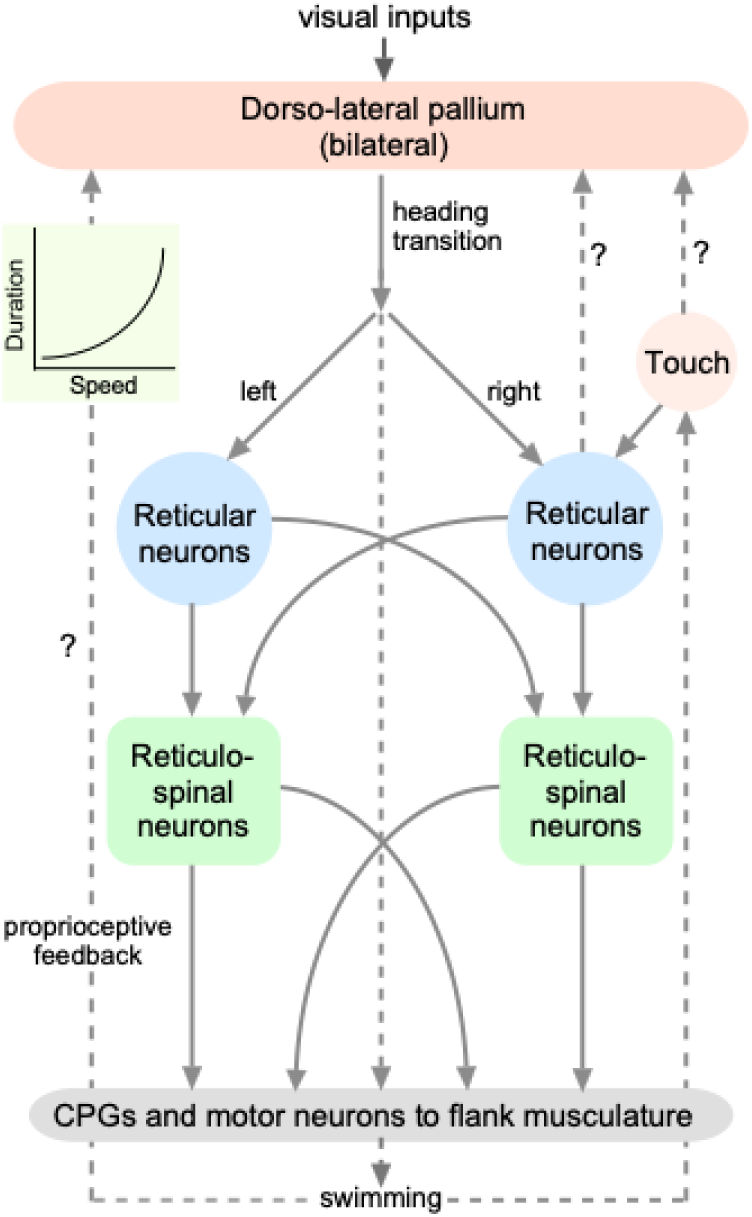
A simplified schematic representation for PI-involved regions and their connections in the fish brain. Visual inputs, normally present in the natural environment reach the dorsolateral pallium via the optic tectum and the preglomular nucleus (not shown). These are replaced by touch inputs to reticular neurons within a go-and-touch strategy following a forward swim coordinated by reticular and reticulospinal neurons. Proprioceptive feedback (shown on the left) to pallial neurons are expected to contribute to the computation of swim parameters, e.g. speed and duration encoding. Touch inputs (shown on the right) are expected to reach the pallium. Dashed arrows reprsent indirect or unknown pathways. CPG: central pattern generator.

The go-and-touch swim strategy analyzed and represented as Markov plots in our results can also be expressed as a behavioral feature-based ethogram taking advantage of DLC-driven tracking and geometric configuration-driven analysis (Fig. 6C). These observations shed light on how zebrafish adapt their navigational strategies to unfamiliar environments, particularly when deprived of external visual landmarks. Their ability to utilize self-generated sensory cues highlights the importance of proprioception and touch in spatial mapping. The STR maneuver emerges as a transitional tactic, enabling zebrafish to iteratively update their map of boundaries while progressively shifting toward more exploratory behaviors.

### 4.5. Neural circuits and mechanisms for path integration

The zebrafish brain provides a behaviorally, neurophysiologically, neuronatomically and genetically tractable bridge between insect and mammalian brains. Interestingly, soon after our first report of path integration in zebrafish (Sanghera and Kanwal, 2021), place cells were reported to exist within the telencephalon of zebrafish larvae (Yang et al., 2024). The bauplan of the vertebrate brain is already well established in teleosts (Lal and Kawakami, 2022; Mueller and Wullimann, 2009; Yamamoto et al., 2007). Fish do not possess a well-developed mammalian-level organization of cortical structures, such as the hippocampus and entorhinal cortex, that are known to enable navigation in mammalian species. This calls into question whether well-organized mammalian-like brain structures are essential for navigation or if the same functionality can be achieved by synaptic and circuit level mechanisms as in the ring attractor within the ellipsoid body in the common fruit fly (Kakaria and de Bivort, 2017; Seelig and Jayaraman, 2015)

A cognitive map is typically created by strategic, free-exploration and probabilistic associations between egothetic and allothetic cues (Bingman and Able, 2002; Ferguson et al., 2019; Liu et al., 2019; Tsoar et al., 2011; Zhu et al., 2013). It allows an animal to localize itself within its immediate space and determine the best trajectory to specific destinations. Our knowledge of the underlying neurobiology for spatial navigation is not yet complete, but remarkable progress has been made on the nature of spatial representation in the mammalian brain in terms of the cell types and receptive fields involved (Etienne et al., 1996; Iggena et al., 2023; Langston et al., 2010; Vinepinsky et al., 2020). In mammals, the key players appear to be place cells in the hippocampus and grid cells in the entorhinal cortex (Gustafson and Daw, 2011; Moser et al., 2008; Solstad et al., 2006; Zhu et al., 2013). Place cells fire maximally at a particular location based on multimodal sensory cues, whereas grid cells compute the vector from the starting to goal location (Burgess and O’Keefe, 2011).

Based on our data and neural recordings by others in different fish species, we construct a working model of brain regions and neural operations involved in mapping space within brains of adult zebrafish (Fig. 7). Recent studies in goldfish show that neurons in the lateral pallium encode swim-related parameters such as head direction, swimming speed, and a swim velocity-vector (Vinepinsky et al., 2020). These motion-sensitive neurons likely contribute to PI by detecting and controlling swim direction and speed, possibly through proprioceptive feedback or sensory information about water flow on the skin. The lateral pallium also contains neurons that can detect boundaries of the home space (Vinepinsky et al., 2020). How they generate a code for this is unknown, but inputs from either reticular neurons or inputs about optic flow via the optic tectum may play an important role (Raudies and Hasselmo, 2012).

Reticular neurons in fish have complex touch-sensitive receptive fields restricted to different areas on the lip region that is densely innervated by touch-sensitive receptors (Kanwal and Finger, 1997). The overlapping touch-receptive fields can be ipsi- or bilateral and are activated by gentle punctate or gliding touch (Kanwal and Finger, 1997). Touch, activated by contact with the tank wall, may serve as a waypoint in the map-making strategy and signal integration of swim parameters. Encoding the order of a series of spatially linked sensory inputs is another requirement for a go-and-touch mapping strategy and fish are known to have the capacity to do so (Burt de Perera, 2004). Finally, motor-sensory integration must be coupled with bouts of sustained attention, likely directed by the habenula, to the animal’s swim-speed and heading to create a neural representation of space (Lecourtier and Kelly, 2005). Neural mechanisms for re-orientations and action selection need to be examined within this context as they have been for social aggression behaviors culminating in winner/loser specification (Okamoto et al., 2021).

### 4.6. Implications and future directions

We report here a novel mapping strategy that combines PI with self-generated sensory cues for mapping boundaries and space. Further investigation of PI in zebrafish can lead to a deeper understanding of the genetic underpinnings and evolutionary origin of neural circuits, such as in the hippocampus and of neurological disorders like representational neglect, affecting spatial navigation and working memory (Guariglia et al., 2005). This is possible because of the rapid development and availability of genetic tools that allow precise manipulation of neural activity and observation of resultant behavioral changes in simple animal models, such as the zebrafish. In GCaMP-labeled transgenic zebrafish, neurons are tagged with fluorescent proteins so that their activity can be visualized using two-photon and related imaging methodologies (Muto and Kawakami, 2016; Muto et al., 2011). This can also facilitate fundamental discoveries on the neuroethological aspects of map generation and navigation. For example, how are receptive fields of place- and grid-like cells created. This may be feasible by studying PI behavior in transgenic lines where specific neural circuits can be labeled and chemically ablated to see their effect on behavior (Mruk et al., 2020).

Several questions remain to be addressed. It may be argued that in our setup, zebrafish could have taken advantage of the overhead camera shape as a visual cue for space mapping. We do not believe that to be the case. The short distance of the relatively large rectangular camera face from the water surface is unlikely to change the subtended angle on the fish retina, rendering it inadequate as a relevant cue. Only a shadow cast by an overhead stimulus looming or sweeping at a particular speed triggers attention and an escape reaction (Mancienne et al., 2021). Zebrafish vision is typically directed in front and below. Finally, information about swimming speed derived from optic flow is mostly restricted to lateral regions of the lower visual field (Alexander et al., 2022). To completely rule out the role of visual cues, however, the current study needs to be replicated in the dark. We have recently completed such a study under infra-red light and observed that zebrafish make the same STR maneuvers during the early phase of freely swimming in a novel tank environment (Shraybman and Kanwal, in preparation).

Finally, to arrive at a computational model of map generation, we need to precisely track turn angles, swim speeds, and contact locations at the tank boundary. We need to test if the wall touchpoints function as olfactory markers, allowing them to return to the same location, or perhaps as “reward”, allowing the fish to associate energy expenditure from swimming with time-segments of movement and thereby encouraging tank exploration. Our primary purpose here was to report the use of PI via the STR maneuver that became feasible using a machine-learning-driven classification of pose and action patterns. Future studies will focus on more detailed analysis of PI to test the contribution of swim speed parameters, vector addition, attention and touch integration for map generation. We do not have physiologic or anatomic evidence yet of the generation or existence of a map or some type of spatial representation of space, though place cells were recently reported in the telencephalon,and head direction cells in the brainstem, in larval zebrafish (Petrucco et al., 2023; Yang et al., 2024).

Together, these findings position zebrafish as a powerful non-mammalian vertebrate model for dissecting the circuit-level and neurogenetic mechanisms that support spatial updating and navigation, and suggest that core computational strategies for path integration may arise from conserved sensorimotor principles rather than specialized cortical architectures. Convergence across taxa underscores the importance of PI for efficient navigation in different environments. We hope neuroethological studies in the future will take advantage of our finding to enable a deeper understanding of circuit-level organization and the nature and role of spatial representations within homologous brain structures in teleosts and their evolution into more complex structures in the mammalian brain (Yamamoto et al., 2007).

### 4.7. Summary and conclusions

To summarize, zebrafish adopt a systematic behavioral strategy involving PI to estimate the shape and size of a target space. Early exploration is dominated by boundary-anchored orientation updates, in which tactile contact scaffolds the integration of self-motion cues through structured sequences of reorientation and corrective turning. As exploration proceeds, these same behavioral states become progressively decoupled from immediate boundary contact, indicating that spatial information derived from earlier interactions is internalized and used to guide navigation. This transition from externally anchored to internally coordinated control mirrors fundamental principles of path integration described across invertebrate and mammalian systems, despite the absence of mammalian cortical navigation circuits in teleosts. We also show that at early stages of development, larval zebrafish do not exhibit STR maneuvers, but STR-like maneuvers begin to emerge in approximately one-month-old juvenile zebrafish.

Circuit-level studies and recording of neural activity during the completion of STR maneuvers in adult zebrafish are needed, but at present these are technically challenging. When feasible, these studies can make a significant contribution towards an understanding of the mechanisms of map creation in a simple brain and how gene expression elaborates and functionalizes the relevant circuitry. We propose that by combining idiothetic information generated from body angles and movements with allothetic (touch) cues, zebrafish create a representation of their home space. This can be equated to the use of heading and waypoints as a map-making strategy. our results demonstrate that adult zebrafish reorganize exploratory behavior in a manner consistent with path integration–like navigation.

## Statement of Ethics

All procedures were reviewed and approved by the Georgetown Animal Care and Use Committee (protocol # 2016-1151).

## Author Contributions

J.S.K. conceived, designed and supervised the study after observation of PI-like behavior in adult zebrafish. B.S. applied machine learning methods for tracking and pose analysis. B.S. and J.S.K. contributed to data acquisition. B.S., P.S. and J.S.K. contributed to data analysis and visualization. B.S. presented preliminary data at a scientific conference. J.S.K. wrote the original draft of the manuscript with contributions from B.S. and P.S. to the Methods and Results sections. All authors reviewed and approved the final version of the manuscript.

## Institutional Review Board Statement

The animal study protocol was approved by the Institutional Review Board of Georgetown University (protocol code 2016-1151 and date of approval: 8 December 2022).

## Data Availability

The data that support the findings of this study will be made available upon request via BioSciences. Further enquiries can be directed to the corresponding author.

## Acknowledgements

We thank the Biomedical Graduation Research Organization at Georgetown University for their support in terms of administration and facilities. We thank the Division of Comparative Medicine, and the Zebrafish Shared Resource directed by Dr. Eric Glasgow for their assistance with routine animal care and maintenance. During the preparation of this manuscript/study, the authors used generative AI (ChatGPT, vers. 5.2) for the purposes of literature search, generation of Python code, refining parts of text and data analysis and interpretation. The authors have reviewed and edited the output and take full responsibility for the content of this publication.

