## Supplemental information for "Adult Zebrafish Engage in Path Integration-like Behavior When Exploring a Novel Environment"

### Supplementary material

#### Methods

The B-SOiD program classified postural features duruing presence in the novel. A violin plot (insets in confusion matrices) indicate that the model achieved 0.88 accuracy (Fig. S1A). Overlap between the predicted and true labels is shown as a confusion matrix (Fig. S1B). Averaging across all nine animals resulted in a drop in model accuracy from 0.88 to 0.70.

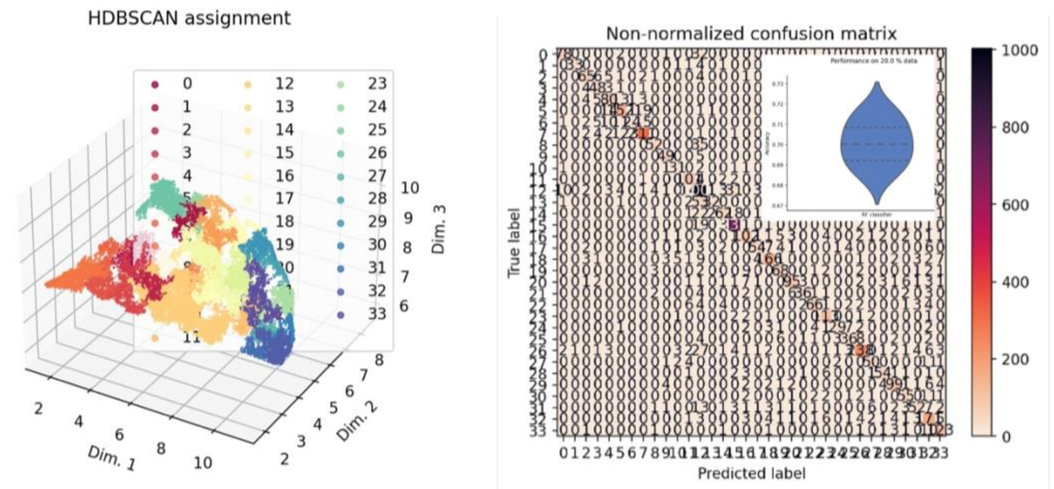


Figure S1. A. Scatterplots showing hierarchical density-based spatial clustering of pose data in the first (left). B-SOiD analysis elucidated 13 distinct behavioral motifs from the first minute and 12 behavioral motifs from the late phase of tracking. Motifs included forward glide and swim, reorienting,sharp turns, and stationary (wait) postures. Using a minimum cluster size of 2.5% yielded ~93% confidence in the classification.

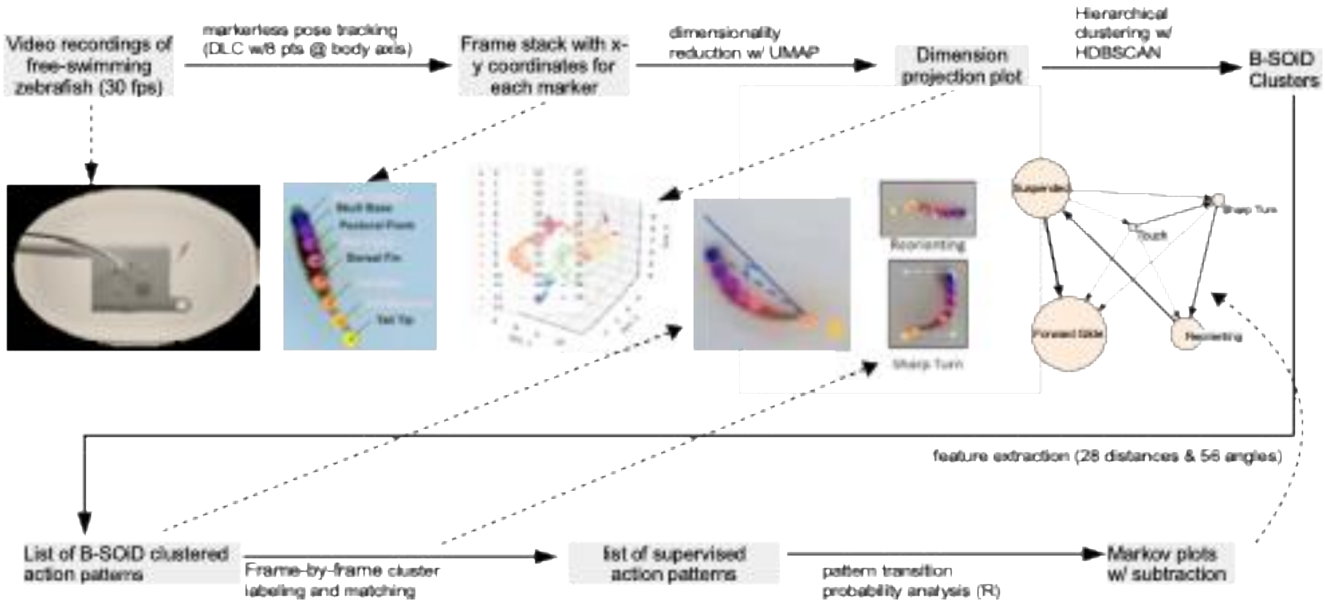

Figure S2. Data acquisition and analysis pipeline to extract postures and swim parameters in adult zebrafish during free exploration in a novel tank environment to establish that fish swim in a non-random pattern.

#### Event Duration and Minima-Based Frequency Measures

Averaged, fish-normalized event compositions revealed that changes in event duration and event initiation frequency were partially dissociable. While duration-based measures emphasized prolonged engagement in specific locomotor states, minima-based analyses captured shifts in the *rate of behavioral switching*.

Difference plots (Minute 8 − Minute 1) demonstrated that some states increased in proportional duration without corresponding increases in initiation frequency, indicating consolidation of behavior rather than increased exploratory sampling. Conversely, reductions in discrete turn initiations occurred even when overall turning occupancy remained stable, consistent with smoother, more sustained trajectories.

To dissociate sustained behavioral engagement from discrete behavioral initiation, two complementary metrics were computed. Duration-based composition, defined as the total number of frames assigned to each event type. Minima-based event frequency, defined as the number of 0→1 onsets for each event, using a minimum separation of 5 frames to avoid over-counting clustered events. For each fish, event counts were normalized to proportions summing to one across all events. Group-level averages were computed across seven fish.

### Results

Difference plots showing changes in average, fish-normalized proportions of behavioral events between Minute 8 and Minute 1. Each fish contributes equally by normalizing event proportions to sum to one prior to averaging. Top: Duration-based composition (event-frame proportions). Bottom: Minima-based event frequency (event initiation rates). Positive values indicate increased relative contribution over time, whereas negative values indicate reductions. Color coding matches Markov network visualizations.

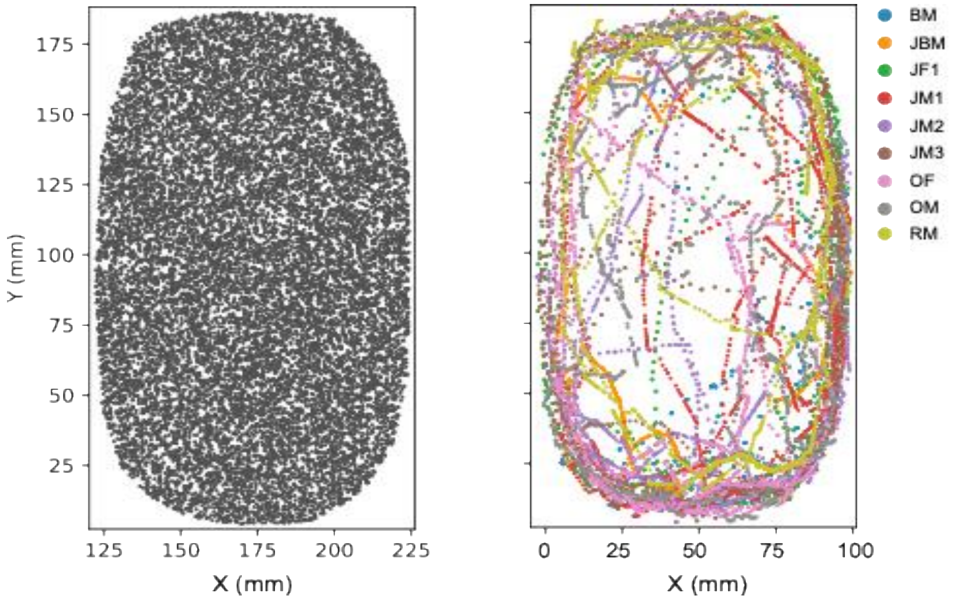


Figure S3. A dot-matrix display of x-y coordinates based on a simulation of random points with smoothing for dimensions matching the test tank. B. A plot of the tracked x-y coordinates of swim trajectories for nine adult zebrafish, identified by color, recorded after their placement in a new tank environment The observed non-random fish exploratory behavior of swimming mostly around the tank parameter from nine animals. The random distribution is approximately even across all distances. Each colored track represents the swim pattern of a different animal during the first minute in a novel tank environment.

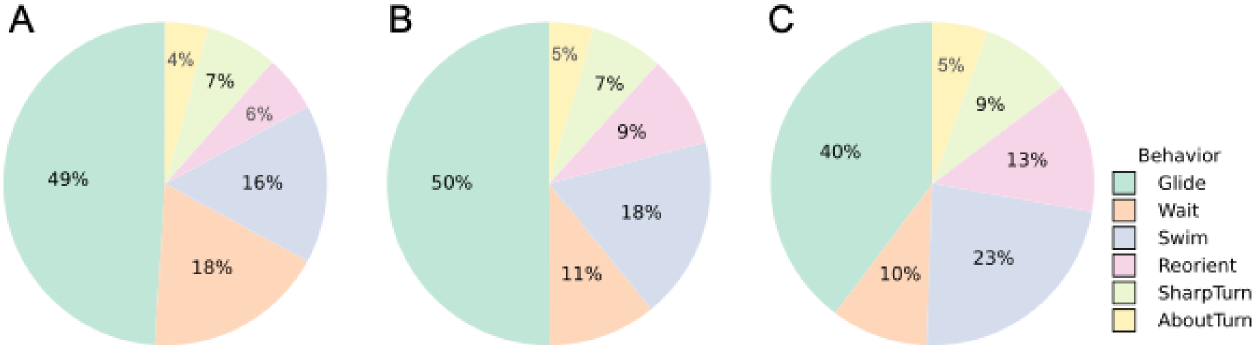


Figure S4. Pie charts showing the changes in proportion of behavioral states during the first, fifthe and tenth minutes of free-swimming and exploration in a novel tank environment.

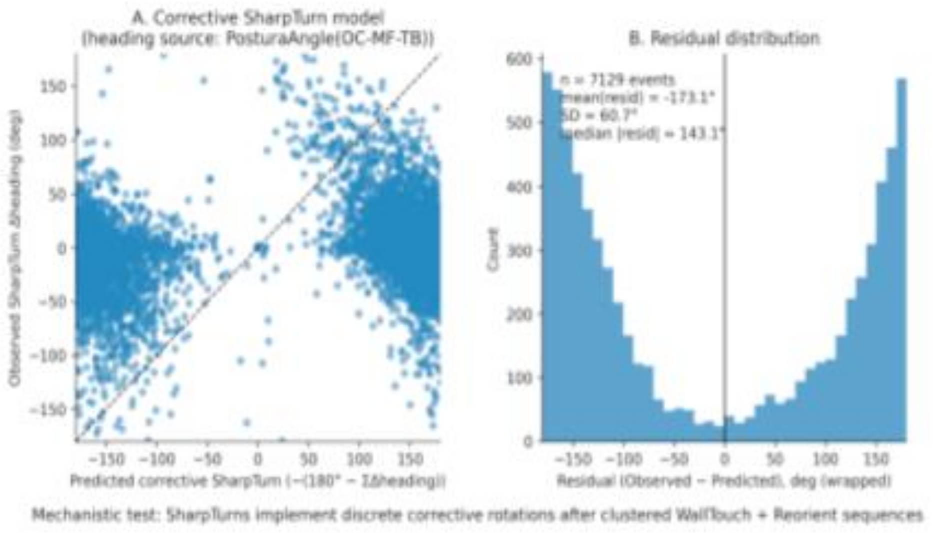


Figure S5. A conceptual schematic summarizing the proposed mechanism: repeated boundary contacts accumulate angular error through discrete heading updates and reorientation, which is compensated by a corrective turn at the third wall encounter.

### Table S1. Paired within-fish statistical comparisons of behavioral states and transitions

Values are means across fish. Minute 1 and Minute 8 are compared using paired Wilcoxon signed-rank tests within individual animals. Significance: ns p≥0.05; * p<0.05; ** p<0.01; *** p<0.001. Transition probabilities are row-normalized within fish.

| Metric | State or Transition | Minute 1 | Minute 8 | Δ (8–1) | Wilcoxon W | p value | Sig. |
|---|---|---|---|---|---|---|---|
| Occupancy | Glide | 0.374 | 0.453 | 0.079 | 6 | 0.219 | ns |
| Occupancy | Reorient | 0.242 | 0.266 | 0.024 | 11 | 0.688 | ns |
| Occupancy | SharpTurn | 0.095 | 0.087 | -0.008 | 11 | 0.688 | ns |
| Occupancy | Swim | 0.080 | 0.091 | 0.010 | 10 | 0.578 | ns |
| Occupancy | Wait | 0.208 | 0.103 | -0.106 | 9 | 0.469 | ns |
| WallTouchOverlap | Glide | 0.108 | 0.216 | 0.108 | 3 | 0.078 | ns |
| WallTouchOverlap | Reorient | 0.126 | 0.201 | 0.075 | 4 | 0.109 | ns |
| WallTouchOverlap | SharpTurn | 0.094 | 0.123 | 0.029 | 4 | 0.345 | ns |
| WallTouchOverlap | Swim | 0.104 | 0.250 | 0.146 | 2 | 0.047 | * |
| WallTouchOverlap | Wait | 0.125 | 0.214 | 0.089 | 5 | 0.156 | ns |
| Transition | Glide → Glide | 0.454 | 0.523 | 0.069 | 9 | 0.469 | ns |
| Transition | Glide → Reorient | 0.241 | 0.249 | 0.008 | 13 | 0.938 | ns |
| Transition | Glide → SharpTurn | 0.059 | 0.053 | -0.006 | 10 | 0.578 | ns |
| Transition | Glide → Swim | 0.111 | 0.079 | -0.031 | 11 | 0.688 | ns |
| Transition | Glide → Wait | 0.136 | 0.095 | -0.040 | 11 | 0.688 | ns |
| Transition | Reorient → Glide | 0.392 | 0.422 | 0.030 | 6 | 0.219 | ns |
